# Connexins evolved after early chordates lost innexin diversity

**DOI:** 10.1101/2021.10.08.463644

**Authors:** Georg Welzel, Stefan Schuster

## Abstract

Gap junction channels are formed by two unrelated protein families. Non-chordates use the primordial innexins, while chordates use connexins that superseded the gap junction function of innexins. Chordates retained innexin-homologs, but N-glycosylation prevents them from forming gap junctions. It is puzzling why chordates seem to exclusively use the new gap junction protein and why no chordates should exist that use non-glycosylated innexins to form gap junctions. Here, we identified glycosylation sites of 2270 innexins from 152 non-chordate and 274 chordate species. Among all chordates, we found not a single innexin without glycosylation sites. Surprisingly, the glycosylation motif is also widespread among non-chordate innexins indicating that glycosylated innexins are not a novelty of chordates. In addition, we discovered a loss of innexin diversity during the early chordate evolution. Most importantly, the most basal living chordates, which lack connexins, exclusively possess innexins with glycosylation sites. A bottleneck effect might thus explain why connexins have become the only protein used to form chordate gap junctions.

## Introduction

Animals from hydra to human use gap junction channels to couple adjacent cells and thus enable direct intercellular communication. Interestingly, gap junction channels are formed by two unrelated integral membrane proteins: innexins and connexins. The innexins are the primordial gap junction proteins that have been identified in all eumetazoans except sponges, placozoa and echinoderms (Slivko-Koltchik et al., 2019). The connexins arose *de novo* during the early chordate evolution and constitute the gap junction channels of all living chordates except lancelets (Abascal et al., 2013; Mikalsen et al., 2021; Slivko-Koltchik et al., 2019). Despite the lack of sequence homology (Alexopoulos et al., 2004), the topology (Maeda et al., 2009; Michalski et al., 2020; Oshima et al., 2016) (Figure 1A, B) and function (Pereda et al., 2017; Skerrett et al., 2017) of connexin- and innexin-based gap junction channels are remarkably similar. Nevertheless, it is thought that chordates have completely replaced the innexin-based gap junctions by the novel connexin-based gap junctions. Vertebrates still express innexin-homologs (Baranova et al., 2004; Panchin et al., 2000), called pannexins, but it is supposed that they stopped forming gap junctions and since then only function as non-junctional membrane channels (Dahl et al., 2014; Esseltine et al., 2016; Sosinsky et al., 2011). This hypothesis is based on the discovery that the three pannexins of humans and mice are glycoproteins. Each of the pannexins contains an identified consensus motif (Asn-X-Ser/Thr) for asparagine (N)-linked glycosylation within either the first or the second extracellular loop (Penuela et al., 2007; Penuela, Simek, et al., 2014; Ruan et al., 2020; Sanchez-Pupo et al., 2018). This enables the posttranslational attachment of sugar moieties at the asparagine residue within the consensus sequence which hinders two pannexin channels of adjacent cells to form intercellular channels (Ruan et al., 2020) (Figure 1C). Based on these findings, it has been assumed that each vertebrate pannexin is equipped with a N-linked glycosylation site (NGS) and thus lost its gap junction function (Sosinsky et al., 2011). However, it remains unclear whether really all vertebrate pannexins are glycosylated and thus presumably only function as single membrane channels. It is also unknown whether glycosylation is indeed a novel modification gained by chordates to prevent their innexin-homologues from forming gap junctions. Specifically, previous studies have shown that at least two non-chordate species, *Aedes aegypti* (Calkins et al., 2015) and *Caenorhabditis elegans* (Kaji et al., 2007), possess an innexin protein with an extracellular NGS that can be glycosylated. These findings raise the intriguing possibility that N-glycosylation might actually be rather common in both chordate and non-chordate innexins, and that N-glycosylation might have played an important role in the evolution of gap junction proteins.

**Figure 1.**
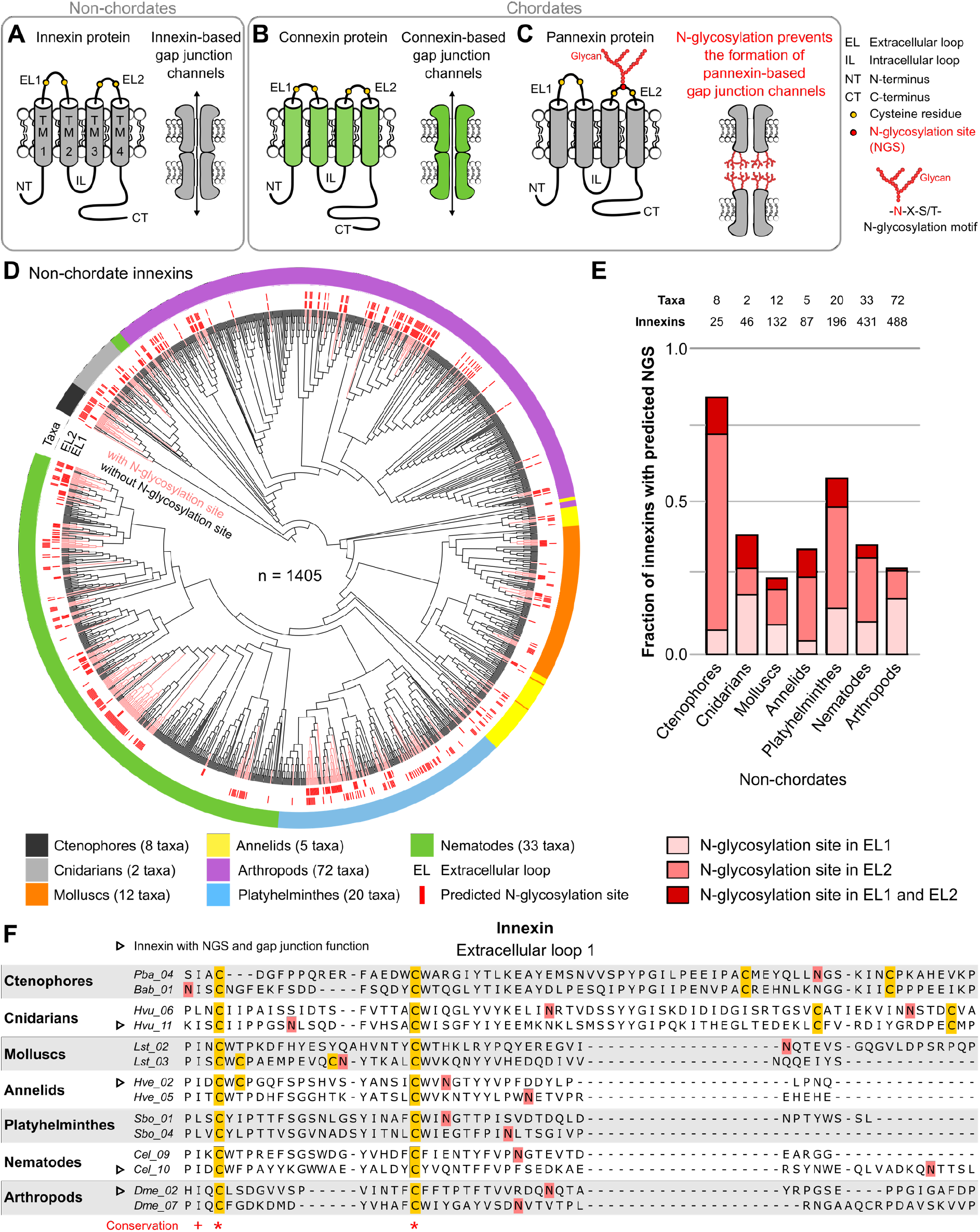
Innexins with N-linked glycosylation sites are widespread among non-chordate animals. (**A**) Innexins and (**B**) connexins are integral membrane proteins that share a common membrane topology with four hydrophobic transmembrane domains (TM) linked by one intracellular (IL) and two extracellular loops (EL). Connexin and innexin proteins can assemble to form a hemichannel. The hemichannels of two neighboring cells can form intercellular gap junction channels that are stabilized by disulfide bonds between cysteine residues in their ELs (Dahl et al., 1991) (yellow circles). (**C**) The innexin homologues of vertebrates, called pannexins, are glycoproteins that contain N-linked glycosylation consensus sites (NGS) within their extracellular loops. The attachment of glycans to NGSs block the disulfide bond formation and hence prevents the proper docking of the hemichannels and the formation of pannexin-based gap junctions (Ruan et al., 2020). (**D**) Maximum likelihood phylogeny of innexin proteins from 152 non-chordate species (n = 1404 innexins). Black and red branches represent innexins without and with NGSs in the extracellular loops, respectively. The red bars in the inner circle indicate whether EL1 or EL2 contains the NGS. The colors of the outer circle represent the phylum classification of the innexins. (**E**) Fraction of innexins with NGSs in their extracellular loops in the taxonomic groups shown in (**D**). (**F**) Representative multiple sequence alignment of the extracellular loop 1 of innexins. The alignment contains selected innexin sequences of representative species of each taxonomic group. The complete alignment including all non-chordate innexins is provided as a supplementary file (Figure 1–source data 2). Conserved residues are highlighted (*, absolutely conserved; +, physicochemical properties are conserved). The cysteine residues (yellow) and the NGSs (red) in each EL1 are colored (note that some innexins have more than two cysteines). The identified NGSs in EL1 and EL2 of the innexins of all other non-chordate species are shown in Figure 1–figure supplement 1. Arrowheads mark innexins with NGSs that have been shown to form functional gap junction channels (Figure 1–source data 3)

Since the experimental identification of N-glycosylated proteins is technically demanding, time consuming and expensive, accurate computational methods are commonly used to identify N-linked glycosylations sites (NGS) in primary amino acid sequences (Gupta et al., 2002; Pitti et al., 2019). In this study, we used the wealth of genomic data that is now available in several public protein and genomic databases to analyze the occurrence of NGSs in non-chordate and chordate innexins in silicio. Based on our findings, we suggest a new evolutionary scenario in which a loss in innexin diversity could explain why the connexins arose *de novo* during the early chordate evolution and why connexins have completely replaced the innexins that so successfully serve diverse functions in the nervous systems of invertebrates.

## Results and Discussion

We first screened for innexin proteins across multiple non-chordate taxa by using innexin proteins as sequence queries in BLAST searches. Only hits that fulfilled defined criteria were included in our study (for more details see Materials and Methods). In total, we collected the amino acid sequences of 1405 non-chordate innexins from 152 species across 7 higher-level taxonomic groups (ctenophores, cnidarians, molluscs, annelids, platyhelminthes, nematodes and arthropods). We subsequently searched in each of the sequences for the consensus motif for N-glycosylation (Asn-X-Ser/Thr). As the extracellular glycosylation of pannexins hinders the gap junction formation (Ruan et al., 2020), we only included NGSs that are located in the extracellular loops of the innexins in our study. Surprisingly, we found that innexins with extracellular NGSs are widespread among the examined non-chordate phyla, comprising more than 80 % of the innexins in the ctenophores (Figure 1D, E, Figure 1–source data 1). The position of the NGSs within the extracellular loops as well as the residues around the N-glycosylation consensus motifs are not conserved between the phyla (Figure 1F). Within the single phyla, we found some innexin orthologs that have highly conserved NGSs and extracellular loops (Figure 1–figure supplement 1). However, we did not find any extracellular NGS that was conserved in all species within a phylum. This finding is presumably based on the phylum-specific diversification of innexins. As shown in previous studies (Abascal et al., 2013; Hasegawa et al., 2014; Moroz et al., 2014), and demonstrated in Figure 1D, innexins originated early in the metazoan evolution and have undergone diversification within the different non-chordate phyla. Thus, innexins with extracellular NGSs evolved independently numerous times within the single phyla.

The wide occurrence of innexins with NGSs in all non-chordate phyla (Figure 1D, E) as well as the experimentally confirmed NGSs (Calkins et al., 2015; Kaji et al., 2007) strongly suggest that a large fraction of non-chordate innexins are glycoproteins. These glycosylated innexin channels might then also not be able to form gap junction channels but rather function as non-junctional channels. However, some of the innexins with identified NGSs have previously been shown to form functional gap junction channels (Figure 1F, Figure 1–source data 3). This means that either the predicted NGSs of these innexins are not glycosylated (Apweiler, 1999) or that glycosylation does not necessarily entail the loss of gap junction function in the diverse innexins of invertebrates.

Our findings demonstrated that non-chordate animals possess a vast diversity of innexins, with and without NGSs, and thus function either as non-junctional membrane channels or as intercellular gap junction channels. This finding is in sharp contrast to the situation in chordates, where all innexins are assumed to be glycosylated and unable to form gap junction channels (Dahl et al., 2014; Esseltine et al., 2016; Sosinsky et al., 2011). But is there really not a single chordate species that uses pannexin-based gap junctions? Up to know, extracellular NGSs were only identified in human (Ruan et al., 2020), mouse (Penuela et al., 2007), rat (Boassa et al., 2007) and zebrafish (Kurtenbach et al., 2013; Prochnow et al., 2009) pannexins. To clarify the prevalence of extracellular NGSs in chordates, we again used public protein and genomic databases to screen for innexin proteins across multiple chordate taxa. In total, we collected the amino acid sequences of 865 chordate innexins from 274 species across 9 higher-level taxonomic groups (lancelets, tunicates, lampreys, cartilaginous fish, bony fish, amphibians, reptiles, birds, and mammals) and then searched, as previously described, in the extracellular loops of each of the sequences for the consensus motif for N-glycosylation (Asn-X-Ser/Thr). Our results clearly show that each single innexin in every chordate species has at least one NGS in its extracellular loops (Figure 2A, B, Figure 2–source data 1). Moreover, we show that in vertebrates the sequence of the extracellular loops as well as the positions of the glycosylation motifs are highly conserved (Figure 2C-F). Among the three pannexins, conservation is particularly high in the extracellular loops of Pannexin-2. Furthermore, conservation is still seen even after the whole-genome duplication in the common ancestor of teleost fishes (Glasauer et al., 2014), an event that generally provides a source of genetic raw material for evolutionary innovation and functional divergence. Still, each single species retained their pannexins with NGSs (Figure 2F and Supplementary File 4). This is remarkable because a single mutation in the N-glycosylation motif might be sufficient to recover the ability of pannexins to form gap junction channels (Ruan et al., 2020). The surprisingly high conservation of the location and the surrounding sequences of NGSs suggests that the pannexins serve essential roles, with correspondingly high stabilizing selective pressures (Abascal et al., 2013).

**Fig. 2.**
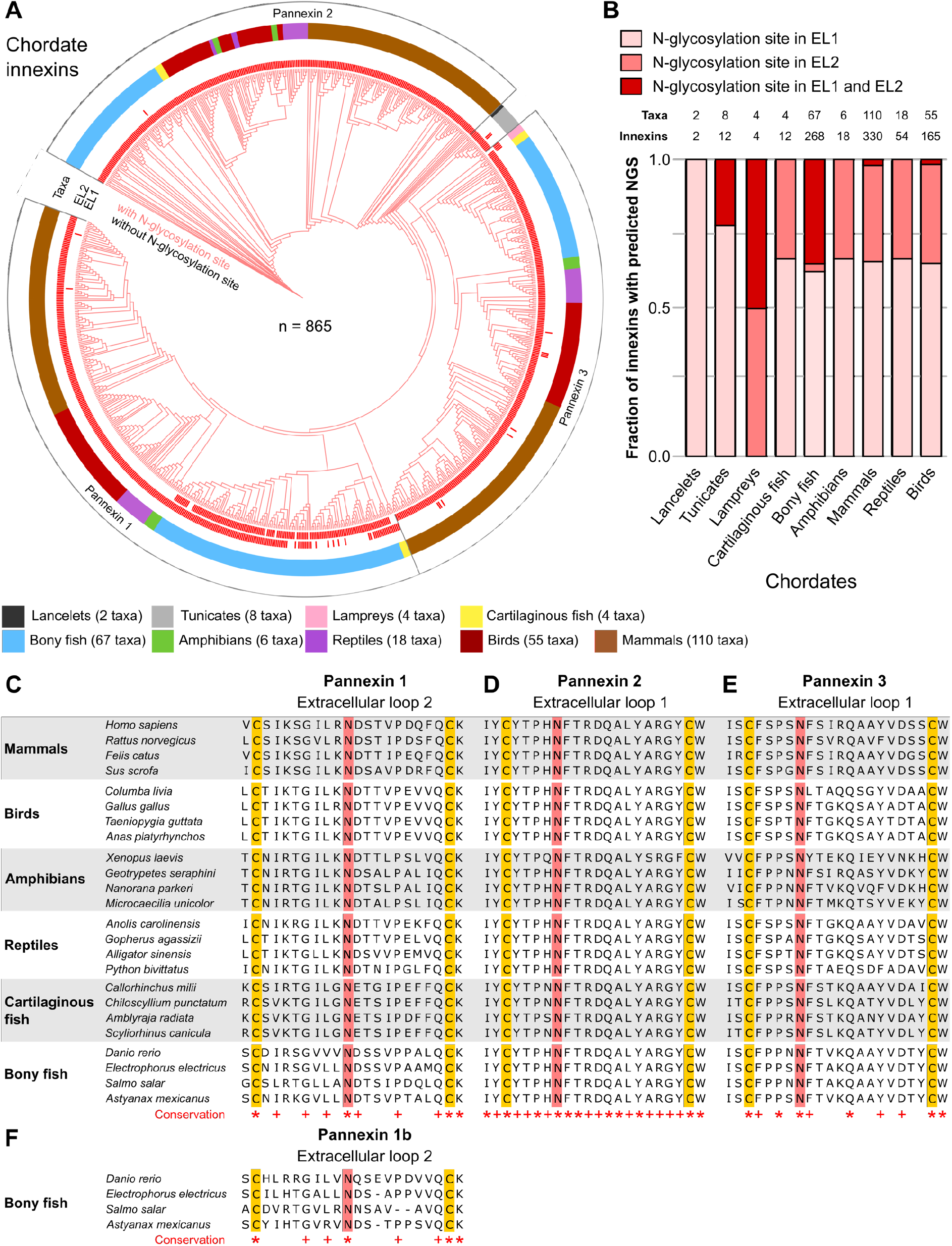
N-linked glycosylation of innexins is highly conserved in chordate animals. (**A**) Maximum likelihood phylogeny of innexin proteins from 274 non-chordate species (n = 865 innexins). Black and red branches represent innexins without and with N-glycosylation consensus sites (NGS) in the extracellular loops, respectively. Note that each innexin in every chordate species has at least one NGS. The red bars in the inner circle indicate whether extracellular loop 1 or 2 (EL1 and EL2) contains the NGS. The colors of the outer circle represent the phylum classification of the innexins. (**B**) Fraction of innexins with NGSs in their extracellular loops (EL) in the taxonomic groups shown in (**A**). (**C**) Representative multiple sequence alignments of the highly conserved extracellular loop 2 of Pannexin 1 and the extracellular loop 1 of (**D**) Pannexin 2 and (**E**) Pannexin 3 in vertebrates. (**F**) Representative multiple sequence alignment of the extracellular loop 2 of the duplicated Pannexin 1 in bony fish. The alignments only contain the sequences of some representative species of each taxonomic group. The complete alignment including all chordate innexins is provided as a supplementary file (Figure 2–source data 2). Conserved residues are highlighted (*, absolutely conserved; +, physicochemical properties are conserved). The cysteine residues (yellow) and the NGSs (red) in each EL are colored. The identified NGSs in EL1 and EL2 of the innexins of all other chordate species are shown in Figure 2–source data 1.

In summary, we show that N-glycosylation is present in both non-chordate and chordate species. Already simple organisms at the beginning of the metazoan evolution attached sugar moieties to some of their innexins to presumably prevent them from forming gap junction channels. In consequence, the vertebrate pannexins did not diverge and change their function driven by the appearance of the connexins but rather originate from an innexin already equipped with NGS. This would be consistent with findings that single membrane channels formed by pannexins and innexins have the same physiological functions and are similar in their biophysical and pharmacological properties (Dahl et al., 2014).

If it is typical for invertebrates to use a great diversity of glycosylated and non-glycosylated innexins and to even form gap junctions from both (Figure 1-figure supplement 2), then the situation in the vertebrates becomes even more puzzling: Why do all vertebrates exclusively retain glycosylated innexins, why do they not form gap junction from them (Ruan et al., 2020) and instead evolved and exclusively use the new connexins for functions that could equally be fulfilled by an innexin? We suggest that looking at the early chordate evolution may solve this puzzle. The chordates are comprised of three subphyla: the lancelets, the tunicates, and the vertebrates. The lancelets represent the most basal chordate lineage that diverged before the split between tunicates and vertebrates (Putnam et al., 2008). The vertebrates split into the jawless fish (lampreys), the most ancient vertebrate group (Smith et al., 2018), and the jawed vertebrates (Figure 3B). As our previous analysis revealed, the lancelets and the lampreys as well as most of the tunicates have only one innexin. In the jawed vertebrates, we find three innexins (called pannexins) in each single species. This is expected from the two whole-genome duplications at the early vertebrate lineage leading first to Pannexin-2 and afterwards to Pannexin-1 and Pannexin-3 (Abascal et al., 2013; Fushiki et al., 2010). Only teleost fishes have a fourth pannexin generated by a whole-genome duplication event during the teleost evolution (Bond et al., 2012; Glasauer et al., 2014). The limited genetic diversity is thus in strong contrast to the rich innexin diversity within the non-chordate phyla (Figure 1C) (Abascal et al., 2013; Hasegawa et al., 2014; Moroz et al., 2014). Moreover, we identified extracellular NGSs in each of the innexins of lancelets, tunicates, and lampreys (Figure 2A, B and Figure 3A). The innexin sequences of these groups are less conserved compared to those of jawed vertebrates and the position of extracellular NGSs are different in lancelets and tunicates. The most important finding is that the sequence of the only innexin of lancelets, which do not yet express connexins (Mikalsen et al., 2021; Slivko-Koltchik et al., 2019) (Figure 3D), contains a NGS in its extracellular loop 1. This suggests that the most basal chordates not only had a limited number of innexins but might also not be able to form functional gap junctions. Interestingly, at this time of the chordate evolution, the connexins arose *de novo* (Figure 1B-D) and developed into diverse gene families (up to 22 connexins in mammals and 46 connexins in bony fish) (Mikalsen et al., 2021).

**Figure 3.**
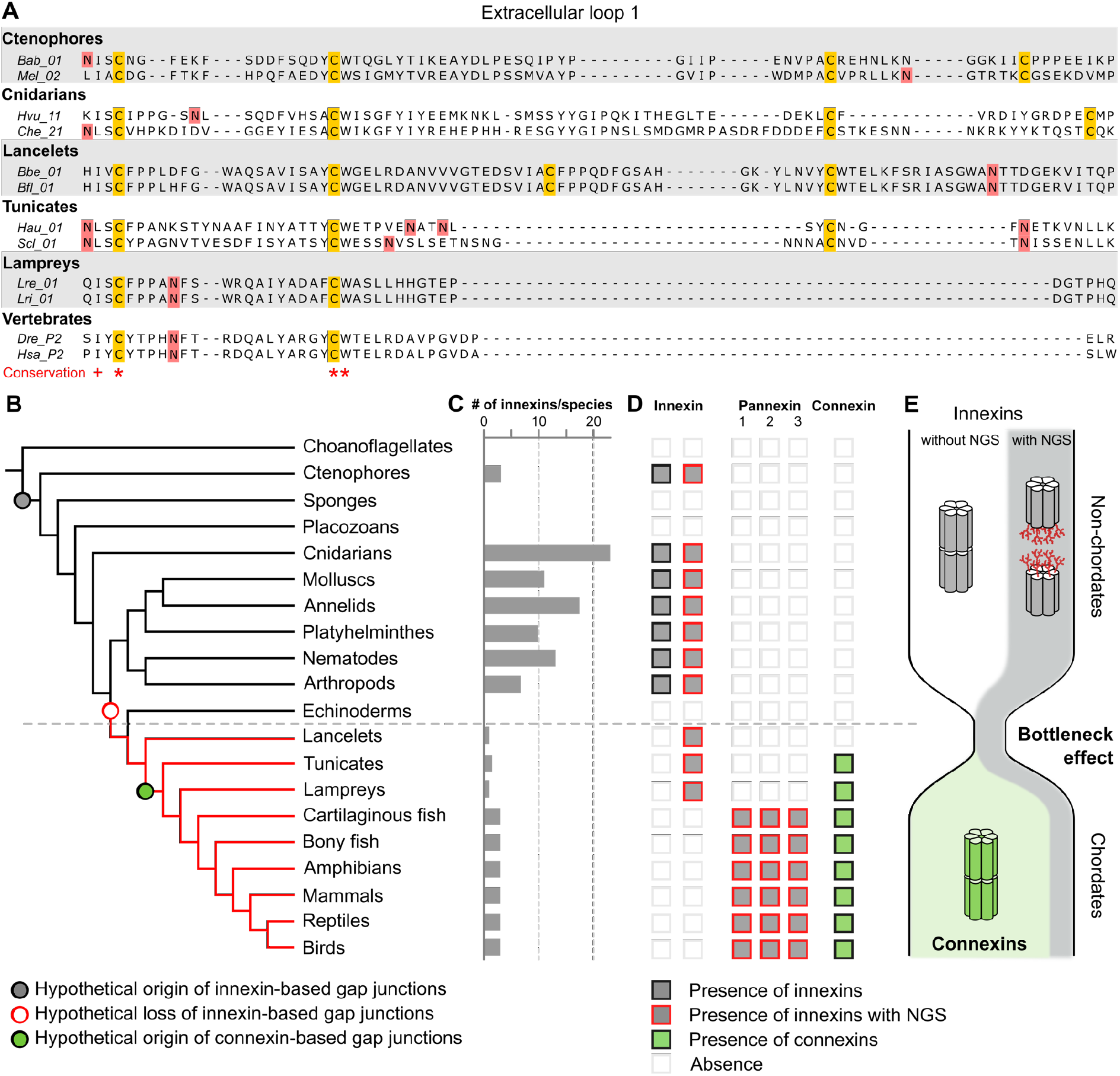
A new evolutionary scenario to explain the origin and exclusive use of connexins in vertebrates. (**A**) Representative multiple sequence alignment of the extracellular loop 1 of chordate and non-chordate innexins. Only extracellular loop 1 is shown as it is one of the best-conserved regions of non-chordate and chordate innexins (Yen et al., 2007). Conserved residues are highlighted (*, absolutely conserved; +, physicochemical properties are conserved). The cysteine residues (yellow) and the NGSs (red) are colored. Note that the lancelets and tunicates have more than two cysteine residues in extracellular loop 1, like cnidarians and ctenophores. (**B**) Simplified phylogenetic tree to visualize the hypothetical loss of innexin-based gap junctions and the origin of connexin-based gap junctions. Lineages that only possess innexins with extracellular NGSs are indicated by a red branch color. (**C**) Mean number of different innexins per species in each of the different phyla. (**D**) Occurrence of innexins and connexins in different metazoans. Innexins with a N-glycosylation consensus site (NGS) in their extracellular domains are present in chordates and non-chordates. Note that lancelets only possess innexins with NGSs and lack connexins. (**E**) A hypothetical evolutionary scenario in which the connexin-based gap junctions evolved after innexin diversity was lost during a bottleneck event in the early chordate evolution.

Based on our findings, we propose that a bottleneck effect at the origin of chordates might have been crucial for the evolution of the novel connexins (Figure 3E). In this evolutionary scenario, innexins were recruited as gap junction proteins in the common cnidarian/bilaterian ancestor. While the innexins functionally diverge in cnidarians and protostomes, the last common ancestor of the deuterostomes had lost all diverse innexins and retained only one that presumably was glycosylated and did not form gap junction channels (Figure 3B). The high conservation of NGSs in the vertebrate innexins that we describe here (Figure 2C-F), their expression in every organ (Penuela et al. 2014) and their association with a variety of diseases (Esseltine et al., 2016; Penuela et al., 2014) suggest that the non-junctional innexin channels already served essential physiological functions in the basal chordates and could not be converted into gap junctions. The loss of innexin diversity on the one hand and the strict conservation of the NGSs in the remaining innexin could thus explain rather simply why the connexin family arose *de novo* and why it became the exclusive gap-junction protein in all deuterostomes although innexin-based gap junctions would have been fully capable to serve all functions (Baker et al., 2014; Bao et al., 2007; Bhattacharya et al., 2019; Lane et al., 2018; Liu et al., 2016; Phelan et al., 2001; Skerrett et al., 2017; Welzel et al., 2018; Yaksi et al., 2010) as they do so successfully in the sophisticated nervous systems of invertebrates (Calabrese et al., 2016; Hall, 2017; Kristan et al., 2005; Marder et al., 2005; Otopalik et al., 2019).

## Materials and Methods

### Database searches

We used public databases to collect innexin amino acid sequences of chordate and non-chordate species. The taxonomic groups that we have analyzed in this study were constrained by the availability of publicly available genomic data. We screened for innexin proteins across multiple taxa by using diverse sequences of the innexin family (PF00876) as sequence queries in BLAST searches. All retrieved sequences were further assessed and only innexin sequences that fulfilled all the following properties were included into our analyses: (1) The sequence was already assigned to the innexin family (PF00876) or a reciprocal BLAST with the sequence hit as query against the UniProt database identified a known innexin sequence as a top hit; (2) The sequence is predicted to contain four transmembrane domains that are connected by two extracellular and one intracellular loop as well as an intracellular N- and C-terminus (see Figure 1A). To clarify this, we used the TMHMM Server v2.0 (http://www.cbs.dtu.dk/services/TMHMM/) to predict membrane topology; (3) The sequence is not fragmented or a duplicate entry. In total, we retrieved 1405 innexin protein sequences of seven non-chordate groups (phylum ctenophores, phylum cnidarians, phylum molluscs, phylum annelids, phylum plathyhelminthes, phylum nematodes and phylum arthropods) and 865 sequences of nine chordate groups (subphylum lancelets, subphylum tunicates, class lampreys, class cartilaginous fish, superclass bony fish, class amphibians, class reptiles, class birds and class mammals). All innexin sequences of molluscs, annelids, plathyhelminthes, nematodes, arthropods, cartilaginous fish, bony fish, amphibians, reptiles, birds, and mammals were obtained from the protein databases at NCBI (http://www.ncbi.nlm.nih.gov) and UniProt (http://www.uniprot.org). The innexin sequences of the ctenophore species were obtained from the Neurobase genome database (http://neurobase.rc.ufl.edu/Pleurobrachia). The innexin sequences of the cnidarian species were obtained from UniProt and the Marimba genome database (http://marimba.obs-vlfr.fr). The LanceletDB database (http://genome.bucm.edu.cn/lancelet) was used to retrieve innexin sequences of lancelets. The innexin sequences of the tunicate species were obtained from NCBI and the ANISEED database (https://www.aniseed.cnrs.fr). The innexin sequences of lampreys were retrieved from the NCBI and the SIMRBASE database (https://genomes.stowers.org). The full list of species and taxa, along with accession numbers and links to the corresponding databases can be found in Figure 1-source data 1 and Figure 2-source data 1.

### Identification of potential N-glycosylation sites (NGS)

To identify potential N-glycosylation sites within the extracellular loops of the non-chordate and chordate innexins, we generated 16 multiple sequence alignments for each taxonomic group (seven non-chordate and nine chordate groups). For each group, we first imported all innexin protein sequences of each species into the Jalview software (version 2.11.1.4) (Waterhouse et al., 2009). The innexin sequences obtained from the UniProt database were automatically retrieved into Jalview by the UniProt sequence fetcher. The sequences obtained from other databases were manually added to Jalview. After aligning the innexin sequences with ClustalW (Thompson et al., 1994), the resulting multiple sequence alignments of each group were used to identify potential N-glycosylation consensus sites (NGS) in the extracellular domains of each innexin protein. NGSs in innexins were predicted by the NetNGlyc 1.0 Server (http://www.cbs.dtu.dk/services/NetNGlyc/) that uses an artificial neural network to examine the sequence context of the N-X-S/T motif. We used the following criteria to include the NGS into our analyses: (1) X in the N-X-S/T motif could be any AA except proline; (2) the potential score was > 0.5 and the agreement between the nine artificial neural networks was ≥ 5/9; (3) the NGS was located in extracellular loop 1 or 2. The positions of all extracellular N-glycosylation sites are reported in Figure 1-source data 1 and Figure 2-source data 1.

### Phylogenomic tree construction

We visualized the incidence of innexins with N-glycosylation motif in their extracellular domains within different taxonomic groups by using phylogenetic trees. To generate a phylogenetic tree of the 1398 non-chordate and the 865 chordate innexins, respectively, we first created two global alignments including the available alignments of the seven non-chordate or the nine chordate groups. Both alignments were generated using MEGA version X (Kumar et al., 2018; Stecher et al., 2020) with the default parameters of ClustalW. Both multiple sequence alignments were then processed by the G-blocks server (http://molevol.cmima.csic.es/castresana/Gblocks.html) (Castresana, 2000) to automatically detect and remove poorly aligned, nonhomologous, and excessively divergent alignment columns. We reconstructed a phylogenetic tree of the non-chordate and the chordate innexins, respectively, by using the raxmlGUI 2.0 software (Edler et al., 2021). Before the phylogenetic analyses, ModelTest-NG (Darriba et al., 2020) was run on the two trimmed alignments with the default parameters to determine the best probabilistic model of sequence evolution. Both phylogenetic trees were built using the maximum likelihood (ML) method based on the JTT model and 100 bootstrap replications. The phylogenetic trees of chordate and non-chordate innexins were visualized, edited and annotated with iTOL v5 (https://itol.embl.de) (Letunic et al., 2021).

## Acknowledgements

We thank Antje Halwas for generating the multiple sequence alignments and Andreas Möglich for inspiring discussions.

## Data availability

All data generated or analyzed during this study are included in the manuscript and in the supporting files.

---

**Figure 1–figure supplement 1.**
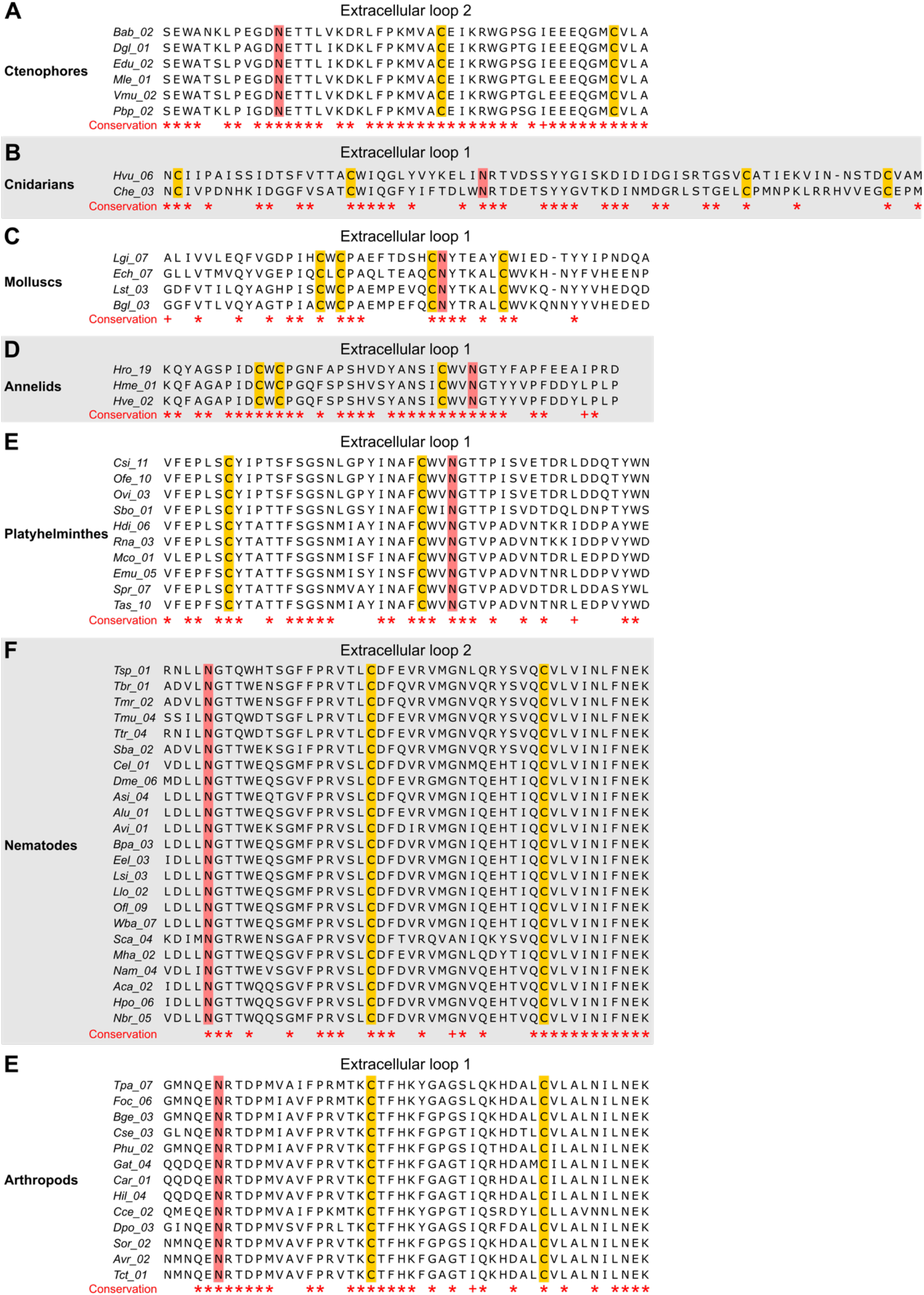
The N-linked glycosylation sites (NGS) in the extracellular loops of innexins are conserved in some orthologs. The multiple sequence alignments show innexin orthologs in (**A**) ctenophores, (**B**) cnidarians, (**C**) molluscs, (**D**) annelids, (**E**) platyhelminthes, (**F**) nematodes and (**E**) arthropods that have highly conserved NGSs in their extracellular loop 1 or 2. Note that we did not find any extracellular NGS that was conserved in all species within a phylum. Conserved residues are highlighted (*, absolutely conserved; +, physicochemical properties are conserved). The cysteine residues (yellow) and the NGSs (red) in each EL are coloured. The identified NGSs in EL1 and EL2 of the innexins of all other non-chordate species are shown in Supplementary File 1.

## References

Abascal, F., & Zardoya, R. (2013). Evolutionary analyses of gap junction protein families. Biochimica et Biophysica Acta (BBA) - Biomembranes, 1828(1), 4–14. doi: 10.1016/j.bbamem.2012.02.007

Alexopoulos, H., Böttger, A., Fischer, S., Levin, A., Wolf, A., Fujisawa, T., Hayakawa, S., Gojobori, T., Davies, J. A., David, C. N., & Bacon, J. P. (2004). Evolution of gap junctions: the missing link? Current Biology, 14(20), R879–R880. doi: 10.1016/j.cub.2004.09.067

Apweiler, R. (1999). On the frequency of protein glycosylation, as deduced from analysis of the SWISS-PROT database. Biochimica et Biophysica Acta (BBA) - General Subjects, 1473(1), 4–8. doi: 10.1016/S0304-4165(99)00165-8

Baker, M. W., & Macagno, E. R. (2014). Control of neuronal morphology and connectivity: Emerging developmental roles for gap junctional proteins. FEBS Letters, 588(8), 1470–1479. doi: 10.1016/j.febslet.2014.02.010

Bao, L., Samuels, S., Locovei, S., Macagno, E. R., Muller, K. J., & Dahl, G. (2007). Innexins form two types of channels. FEBS Letters, 581(29), 5703–5708. doi: 10.1016/j.febslet.2007.11.030

Baranova, A., Ivanov, D., Petrash, N., Pestova, A., Skoblov, M., Kelmanson, I., Shagin, D., Nazarenko, S., Geraymovych, E., Litvin, O., Tiunova, A., Born, T. L., Usman, N., Staroverov, D., Lukyanov, S., & Panchin, Y. (2004). The mammalian pannexin family is homologous to the invertebrate innexin gap junction proteins. Genomics, 83(4), 706–716. doi: 10.1016/j.ygeno.2003.09.025

Bhattacharya, A., Aghayeva, U., Berghoff, E. G., & Hobert, O. (2019). Plasticity of the electrical connectome of C. elegans. Cell, 176(5), 1174–1189.e16. doi: 10.1016/j.cell.2018.12.024

Boassa, D., Ambrosi, C., Qiu, F., Dahl, G., Gaietta, G., & Sosinsky, G. (2007). Pannexin1 channels contain a glycosylation site that targets the hexamer to the plasma membrane. Journal of Biological Chemistry, 282(43), 31733–31743. doi: 10.1074/jbc.M702422200

Bond, S. R., Wang, N., Leybaert, L., & Naus, C. C. (2012). Pannexin 1 ohnologs in the teleost lineage. The Journal of Membrane Biology, 245(8), 483–493. doi: 10.1007/s00232-012-9497-4

Calabrese, R. L., Norris, B. J., & Wenning, A. (2016). The neural control of heartbeat in invertebrates. Current Opinion in Neurobiology, 41, 68–77. doi: 10.1016/j.conb.2016.08.004

Calkins, T. L., Woods-Acevedo, M. A., Hildebrandt, O., & Piermarini, P. M. (2015). The molecular and immunochemical expression of innexins in the yellow fever mosquito, Aedes aegypti: Insights into putative life stage-and tissue-specific functions of gap junctions. Comparative Biochemistry and Physiology Part B: Biochemistry and Molecular Biology, 183, 11–21. doi: 10.1016/j.cbpb.2014.11.013

Castresana, J. (2000). Selection of conserved blocks from multiple alignments for their use in phylogenetic analysis. Molecular Biology and Evolution, 17(4), 540–552. doi: 10.1093/oxfordjournals.molbev.a026334

Dahl, G., Levine, E., Rabadan-Diehl, C., & Werner, R. (1991). Cell/cell channel formation involves disulfide exchange. European Journal of Biochemistry, 197(1), 141–144. doi: 10.1111/j.1432-1033.1991.tb15892.x

Dahl, G., & Muller, K. J. (2014). Innexin and pannexin channels and their signaling. FEBS Letters, 588(8), 1396–1402. doi: 10.1016/j.febslet.2014.03.007

Darriba, Di., Posada, D., Kozlov, A. M., Stamatakis, A., Morel, B., & Flouri, T. (2020). ModelTest-NG: A new and scalable tool for the selection of DNA and protein evolutionary models. Molecular Biology and Evolution, 37(1), 291–294. doi: 10.1093/molbev/msz189

Edler, D., Klein, J., Antonelli, A., & Silvestro, D. (2021). raxmlGUI 2.0: A graphical interface and toolkit for phylogenetic analyses using RAxML. Methods in Ecology and Evolution, 12(2), 373–377. doi: 10.1111/2041-210X.13512

Esseltine, J. L., & Laird, D. W. (2016). Next-generation connexin and pannexin cell biology. Trends in Cell Biology, 26(12), 944–955. doi: 10.1016/j.tcb.2016.06.003

Fushiki, D., Hamada, Y., Yoshimura, R., & Endo, Y. (2010). Phylogenetic and bioinformatic analysis of gap junction-related proteins, innexins, pannexins and connexins. Biomedical Research, 31(2), 133–142. doi: 10.2220/biomedres.31.133

Glasauer, S. M. K., & Neuhauss, S. C. F. (2014). Whole-genome duplication in teleost fishes and its evolutionary consequences. Molecular Genetics and Genomics, 289(6), 1045–1060. doi: 10.1007/s00438-014-0889-2

Gupta, R., & Brunak, S. (2002). Prediction of glycosylation across the human proteome and the correlation to protein function. Pacific Symposium on Biocomputing. Pacific Symposium on Biocomputing, 310–322. doi: 10.1142/9789812799623_0029

Hall, D. H. (2017). Gap junctions in C. elegans : Their roles in behavior and development. Developmental Neurobiology, 77(5), 587–596. doi: 10.1002/dneu.22408

Hasegawa, D. K., & Turnbull, M. W. (2014). Recent findings in evolution and function of insect innexins. FEBS Letters, 588(8), 1403–1410. doi: 10.1016/j.febslet.2014.03.006

Kaji, H., Kamiie, J., Kawakami, H., Kido, K., Yamauchi, Y., Shinkawa, T., Taoka, M., Takahashi, N., & Isobe, T. (2007). Proteomics reveals N-linked glycoprotein diversity in Caenorhabditis elegans and suggests an atypical translocation mechanism for integral membrane proteins. Molecular & Cellular Proteomics, 6(12), 2100–2109. doi: 10.1074/mcp.M600392-MCP200

Kristan, W. B., Calabrese, R. L., & Friesen, W. O. (2005). Neuronal control of leech behavior. Progress in Neurobiology, 76(5), 279–327. doi: 10.1016/j.pneurobio.2005.09.004

Kumar, S., Stecher, G., Li, M., Knyaz, C., & Tamura, K. (2018). MEGA X: Molecular evolutionary genetics analysis across computing platforms. Molecular Biology and Evolution, 35(6), 1547–1549. doi: 10.1093/molbev/msy096

Kurtenbach, S., Prochnow, N., Kurtenbach, S., Klooster, J., Zoidl, C., Dermietzel, R., Kamermans, M., & Zoidl, G. (2013). Pannexin1 channel proteins in the zebrafish retina have shared and unique properties. PLoS ONE, 8(10), e77722. doi: 10.1371/journal.pone.0077722

Lane, B. J., Kick, D. R., Wilson, D. K., Nair, S. S., & Schulz, D. J. (2018). Dopamine maintains network synchrony via direct modulation of gap junctions in the crustacean cardiac ganglion. ELife, 7. doi: 10.7554/eLife.39368

Letunic, I., & Bork, P. (2021). Interactive Tree Of Life (iTOL) v5: an online tool for phylogenetic tree display and annotation. Nucleic Acids Research, 49(W1), W293–W296. doi: 10.1093/nar/gkab301

Liu, Q., Yang, X., Tian, J., Gao, Z., Wang, M., Li, Y., & Guo, A. (2016). Gap junction networks in mushroom bodies participate in visual learning and memory in Drosophila. ELife, 5. doi: 10.7554/eLife.13238

Maeda, S., Nakagawa, S., Suga, M., Yamashita, E., Oshima, A., Fujiyoshi, Y., & Tsukihara, T. (2009). Structure of the connexin 26 gap junction channel at 3.5 Å resolution. Nature, 458(7238), 597–602. doi: 10.1038/nature07869

Marder, E., Bucher, D., Schulz, D. J., & Taylor, A. L. (2005). Invertebrate central pattern generation moves along. Current Biology, 15(17), R685–R699. doi: 10.1016/j.cub.2005.08.022

Michalski, K., Syrjanen, J. L., Henze, E., Kumpf, J., Furukawa, H., & Kawate, T. (2020). The Cryo-EM structure of pannexin 1 reveals unique motifs for ion selection and inhibition. ELife, 9. doi: 10.7554/eLife.54670

Mikalsen, S.-O., Kongsstovu, S. í, & Tausen, M. (2021). Connexins during 500 million years—from cyclostomes to mammals. International Journal of Molecular Sciences 2021, Vol. 22, page 1584, 22(4), 1584. doi: 10.3390/IJMS22041584

Moroz, L. L., Kocot, K. M., Citarella, M. R., Dosung, S., Norekian, T. P., Povolotskaya, I. S., Grigorenko, A. P., Dailey, C., Berezikov, E., Buckley, K. M., Ptitsyn, A., Reshetov, D., Mukherjee, K., Moroz, T. P., Bobkova, Y., Yu, F., Kapitonov, V. V., Jurka, J., Bobkov, Y. V., … Kohn, A. B. (2014). The ctenophore genome and the evolutionary origins of neural systems. Nature, 510(7503), 109–114. doi: 10.1038/nature13400

Oshima, A., Tani, K., & Fujiyoshi, Y. (2016). Atomic structure of the innexin-6 gap junction channel determined by cryo-EM. Nature Communications, 7, 13681. doi: 10.1038/ncomms13681

Otopalik, A. G., Lane, B., Schulz, D. J., & Marder, E. (2019). Innexin expression in electrically coupled motor circuits. Neuroscience Letters, 695, 19–24. doi: 10.1016/j.neulet.2017.07.016

Panchin, Y., Kelmanson, I., Matz, M., Lukyanov, K., Usman, N., & Lukyanov, S. (2000). A ubiquitous family of putative gap junction molecules. Current Biology, 10(13), R473–R474. doi: 10.1016/S0960-9822(00)00576-5

Penuela, S., Bhalla, R., Gong, X.-Q., Cowan, K. N., Celetti, S. J., Cowan, B. J., Bai, D., Shao, Q., & Laird, D. W. (2007). Pannexin 1 and pannexin 3 are glycoproteins that exhibit many distinct characteristics from the connexin family of gap junction proteins. Journal of Cell Science, 120(21), 3772–3783. doi: 10.1242/jcs.009514

Penuela, S., Harland, L., Simek, J., & Laird, D. W. (2014). Pannexin channels and their links to human disease. Biochemical Journal, 461(3), 371–381. doi: 10.1042/BJ20140447

Penuela, S., Simek, J., & Thompson, R. J. (2014). Regulation of pannexin channels by post-translational modifications. FEBS Letters, 588(8), 1411–1415. doi: 10.1016/j.febslet.2014.01.028

Pereda, A. E., & Macagno, E. (2017). Electrical transmission: Two structures, same functions? Developmental Neurobiology, 77(5), 517–521. doi: 10.1002/dneu.22488

Phelan, P., & Starich, T. A. (2001). Innexins get into the gap. BioEssays, 23(5), 388–396. doi: 10.1002/bies.1057

Pitti, T., Chen, C.-T., Lin, H.-N., Choong, W.-K., Hsu, W.-L., & Sung, T.-Y. (2019). N-GlyDE: a two-stage N-linked glycosylation site prediction incorporating gapped dipeptides and pattern-based encoding. Scientific Reports, 9(1), 15975. doi: 10.1038/s41598-019-52341-z

Prochnow, N., Hoffmann, S., Vroman, R., Klooster, J., Bunse, S., Kamermans, M., Dermietzel, R., & Zoidl, G. (2009). Pannexin1 in the outer retina of the zebrafish, Danio rerio. Neuroscience, 162(4), 1039–1054. doi: 10.1016/j.neuroscience.2009.04.064

Putnam, N. H., Butts, T., Ferrier, D. E. K., Furlong, R. F., Hellsten, U., Kawashima, T., Robinson-Rechavi, M., Shoguchi, E., Terry, A., Yu, J.-K., Benito-Gutiérrez, E., Dubchak, I., Garcia-Fernàndez, J., Gibson-Brown, J. J., Grigoriev, I. V., Horton, A. C., de Jong, P. J., Jurka, J., Kapitonov, V. V., … Rokhsar, D. S. (2008). The amphioxus genome and the evolution of the chordate karyotype. Nature, 453(7198), 1064–1071. doi: 10.1038/nature06967

Ruan, Z., Orozco, I. J., Du, J., & Lü, W. (2020). Structures of human pannexin 1 reveal ion pathways and mechanism of gating. Nature, 584(7822), 646–651. doi: 10.1038/s41586-020-2357-y

Sanchez-Pupo, R., Johnston, D., & Penuela, S. (2018). N-glycosylation regulates pannexin 2 localization but is not required for interacting with pannexin 1. International Journal of Molecular Sciences, 19(7), 1837. doi: 10.3390/ijms19071837

Skerrett, I. M., & Williams, J. B. (2017). A structural and functional comparison of gap junction channels composed of connexins and innexins. Developmental Neurobiology, 77(5), 522–547. doi: 10.1002/dneu.22447

Slivko-Koltchik, G. A., Kuznetsov, V. P., & Panchin, Y. V. (2019). Are there gap junctions without connexins or pannexins? BMC Evolutionary Biology, 19(1), 5–12. doi: 10.1186/s12862-019-1369-4

Smith, J. J., Timoshevskaya, N., Ye, C., Holt, C., Keinath, M. C., Parker, H. J., Cook, M. E., Hess, J. E., Narum, S. R., Lamanna, F., Kaessmann, H., Timoshevskiy, V. A., Waterbury, C. K. M., Saraceno, C., Wiedemann, L. M., Robb, S. M. C., Baker, C., Eichler, E. E., Hockman, D., … Amemiya, C. T. (2018). The sea lamprey germline genome provides insights into programmed genome rearrangement and vertebrate evolution. Nature Genetics, 50(2), 270–277. doi: 10.1038/s41588-017-0036-1

Sosinsky, G. E., Boassa, D., Dermietzel, R., Duffy, H. S., Laird, D. W., MacVicar, B., Naus, C. C., Penuela, S., Scemes, E., Spray, D. C., Thompson, R. J., Zhao, H.-B., & Dahl, G. (2011). Pannexin channels are not gap junction hemichannels. Channels, 5(3), 193–197. doi: 10.4161/chan.5.3.15765

Stecher, G., Tamura, K., & Kumar, S. (2020). Molecular evolutionary genetics analysis (MEGA) for macOS. Molecular Biology and Evolution, 37(4), 1237–1239. doi: 10.1093/molbev/msz312

Thompson, J. D., Higgins, D. G., & Gibson, T. J. (1994). CLUSTAL W: Improving the sensitivity of progressive multiple sequence alignment through sequence weighting, position-specific gap penalties and weight matrix choice. Nucleic Acids Research, 22(22), 4673–4680. doi: 10.1093/nar/22.22.4673

Waterhouse, A. M., Procter, J. B., Martin, D. M. A., Clamp, M., & Barton, G. J. (2009). Jalview Version 2-A multiple sequence alignment editor and analysis workbench. Bioinformatics, 25(9), 1189–1191. doi: 10.1093/bioinformatics/btp033

Welzel, G., & Schuster, S. (2018). Long-term potentiation in an innexin-based electrical synapse. Scientific Reports, 8(1), 12579. doi: 10.1038/s41598-018-30966-w

Yaksi, E., & Wilson, R. I. (2010). Electrical coupling between olfactory glomeruli. Neuron, 67(6), 1034–1047. doi: 10.1016/j.neuron.2010.08.041

Yen, M. R., & Saier, M. H. (2007). Gap junctional proteins of animals: The innexin/pannexin superfamily. Progress in Biophysics and Molecular Biology, 94(1–2), 5–14. doi: 10.1016/j.pbiomolbio.2007.03.006

